# *Aedes albopictus* is not an arbovirus aficionado – Impacts of sylvatic flavivirus infection in vectors and hosts on mosquito engorgement on non-human primates

**DOI:** 10.1101/2024.02.19.580944

**Authors:** Hélène Cecilia, Benjamin M. Althouse, Sasha R. Azar, Brett A. Moehn, Ruimei Yun, Shannan L. Rossi, Nikos Vasilakis, Kathryn A. Hanley

## Abstract

The contact structure between vertebrate hosts and arthropod vectors plays a key role in the spread of arthropod-borne viruses (arboviruses); thus, it is important to determine whether arbovirus infection of either host or vector alters vector feeding behavior. Here we leveraged a study of the replication dynamics of two arboviruses isolated from their ancestral cycles in paleotropical forests, sylvatic dengue-2 (DENV-2) and Zika (ZIKV), in one non-human primate (NHP) species from the paleotropics (cynomolgus macaques, *Macaca fascicularis*) and one from the neotropics (squirrel monkeys, *Saimiri boliviensis*) to test the effect of both vector and host infection with each virus on completion of blood feeding (engorgement) of the mosquito *Aedes albopictus*. Although mosquitoes were starved and given no choice of hosts, engorgement rates varied dramatically, from 0% to 100%. While neither vector nor host infection systematically affected engorgement, NHP species and body temperature at the time of feeding did. We also interrogated the effect of repeated mosquito bites on cytokine expression and found that epidermal growth factor (EGF) and macrophage migration inhibitory factor (MIF) concentrations were dynamically associated with exposure to mosquito bites. This study highlights the importance of incorporating individual-level heterogeneity of vector biting in arbovirus transmission models.

## Introduction

Transmission dynamics of vector-borne pathogens are shaped by the contact structure between their arthropod vectors and their vertebrate hosts, which in turn depends upon vector attraction to individual hosts [1]. The most important vector-borne pathogens in the context of global human health are those transmitted by mosquitoes, such as *Plasmodium* spp., dengue (DENV) and Zika (ZIKV) virus [2]. Mosquitoes initially detect hosts at a distance through sensory cues such as carbon dioxide (CO_2_) [3–5]. Once stimulated by CO_2_, mosquitoes will target visual cues and odor plumes to find hosts [6, 7]. Ultimately, mosquitoes will make feeding decisions based on more proximal host cues, such as body heat, humidity, and skin odor, which is a product of volatile chemicals released by the skin microbiome [4, 6, 8].

An effect of infection of host or vector on the feeding behavior of mosquitoes would have significant ramifications for patterns of pathogen transmission. To date, most studies testing the impact of infection on mosquito feeding have used *Plasmodium* spp., the parasite that causes malaria [9]. As reviewed by Sanford and Shutler [10] and Busula *et al.* [11], most studies found that host infection with *Plasmodium* enhanced their attractiveness to mosquitoes. Switching perspective to the vector, laboratory studies have shown that *Plasmodium* infection changes mosquito feeding behavior, but such changes depend on the developmental stage of the parasite and *Plasmodium* species tested, among other factors [10, 12]. For both the host and the vector, it remains unclear whether or how these laboratory findings translate to the field, and whether these observations reveal active manipulation by *Plasmodium* to enhance parasite fitness or simply a non-adaptive by-product of infection [10–12].

In contrast to the rich body of work on *Plasmodium* spp., relatively few studies have investigated the effect of infection with arthropod-borne viruses (arboviruses) on mosquito feeding behavior. A review by Cozzarolo *et al.* [13] showed mixed evidence for an impact of host infection on mosquito feeding. On one hand, several species of *Culex* mosquitoes did not differ in their attraction to house sparrows infected with either St. Louis encephalitis virus or Western equine encephalitis virus relative to uninfected house sparrows, and *Aedes taeniorhynchus* were not more attracted to sheep infected with Rift Valley fever virus (RVFV) than uninfected sheep. On the other hand, *Culex annulirostris* were more attracted to chickens infected with Sindbis virus than uninfected chickens, and *Culex pipiens* were more attracted to sheep infected with RVFV. Recently, Zhang et al. [14] reported that mice and humans infected with DENV-2 or ZIKV produced host cues that were more attractive for *Aedes aegypti* and *Ae. albopictus*, the major vectors of both viruses, than their uninfected counterparts.

Similar to studies of host infection, current evidence is inconclusive as to whether arbovirus infection of mosquitoes impacts their feeding behavior, as reviewed by Maire et al. [15]. Several studies have investigated the impact of DENV infection of *Ae. aegypti* on their tendency to feed on mice [16–20], or guinea pigs [21]. DENV-infected mosquitoes probed for longer periods of time than control mosquitoes in some studies [16, 20] but not others [17, 21], showed a longer total feeding time (time to probe and engorge) in some studies [16, 18] but not others [21], and were less likely to take a blood meal in some studies [19] but not others [20]. To our knowledge, only one study [20] has tested the impact of DENV infection on probing efficiency; it showed DENV-infected *Ae. aegypti* probed more often than uninfected mosquitoes to achieve blood satiety. Studies on ZIKV so far only focused on the impact it might have on *Ae. aegypti* host-seeking behavior, precisely its flight activity [15].

A major limitation to most of the existing DENV and ZIKV studies is their reliance on rodents, which are not natural hosts for either virus. DENV and ZIKV originated in sylvatic cycles in zoonotic reservoir hosts, including non-human primates (NHPs) [22, 23] and arboreal *Aedes* mosquitoes in Asia and Africa, respectively [22, 24]. Both spilled over into humans and established human-endemic cycles transmitted by *Ae. aegypti* and *Ae. albopictus* [25]. *Ae. albopictus* may also be involved in spillover and spillback of both viruses [26–28].

Here, we leveraged a study of sylvatic DENV-2 and ZIKV replication dynamics in both paleotropical (cynomolgus macaques, *Macaca fascicularis*) and neotropical (squirrel monkeys, *Saimiri boliviensis*) NHP hosts to test the effect of both host and vector infection with each virus on vector feeding. Cynomolgus macaques are known to become naturally infected with both viruses [23, 29], whereas neither virus has yet been detected in squirrel monkeys. In this study, batches of *Ae. albopictus* were infected via intrathoracic inoculation, grouped into cartons, placed upon the ear of uninfected NHPs and given the opportunity to feed. Then, batches of uninfected *Ae. albopictus* were placed upon the ear of infected monkeys at regular intervals and allowed to feed. In the control arm of the experiment, uninfected *Ae. albopictus* were fed upon each species of monkey and, subsequently, uninfected mosquitoes were fed on control monkeys at the same intervals as for infected monkeys. In every case, engorged and unengorged mosquitoes were counted from every carton, enabling us to test the impact of infection on engorgement. We tested the effect of mosquito infection status (infected or control) when feeding on naïve hosts, as well as the effect of NHP infection status on naïve mosquitoes. We found that arbovirus infection of vectors and hosts does not systematically impact mosquito engorgement, but when it does, it tends to decrease *Ae. albopictus* tendency to engorge. Of the other biological and experimental variables that we analyzed, only NHP species and body temperature at the time of feeding influenced *Ae. albopictus* tendency to engorge. We also identified cytokines, including epidermal growth factor (EGF) and macrophage migration inhibitory factor (MIF), that were dynamically affected by the repeated exposure to uninfected mosquito bites.

## Material and Methods

### Overview

The experiments that generated the data analyzed here have been described in detail in Hanley *et al.* 2023 [30]. Briefly, fifteen (9 females, 7 males) Mauritius origin cynomolgus macaques (*Macaca fascicularis*), between 2.2 and 4.0 years of age and weighing between 2.7 and 5.2 kilograms (kg) at the time of the study, were purchased from Worldwide Primates, Inc (Miami, FL, USA). Twenty-four (12 females, 12 males) squirrel monkeys (*Saimiri boliviensis boliviensis*) between 4 and 14 years of age and weighing between 0.62 kg and 0.89 kg at the time of the study, were purchased from the MD Anderson Center (Bastrop, TX, USA). All NHPs were anesthetized with ketamine before every procedure. The experiments conducted here were approved via UTMB Institutional Animal Care and Use Committee (IACUC) protocol 1912100. Figure 1 provides an overview of the experimental design.

**Figure 1:**
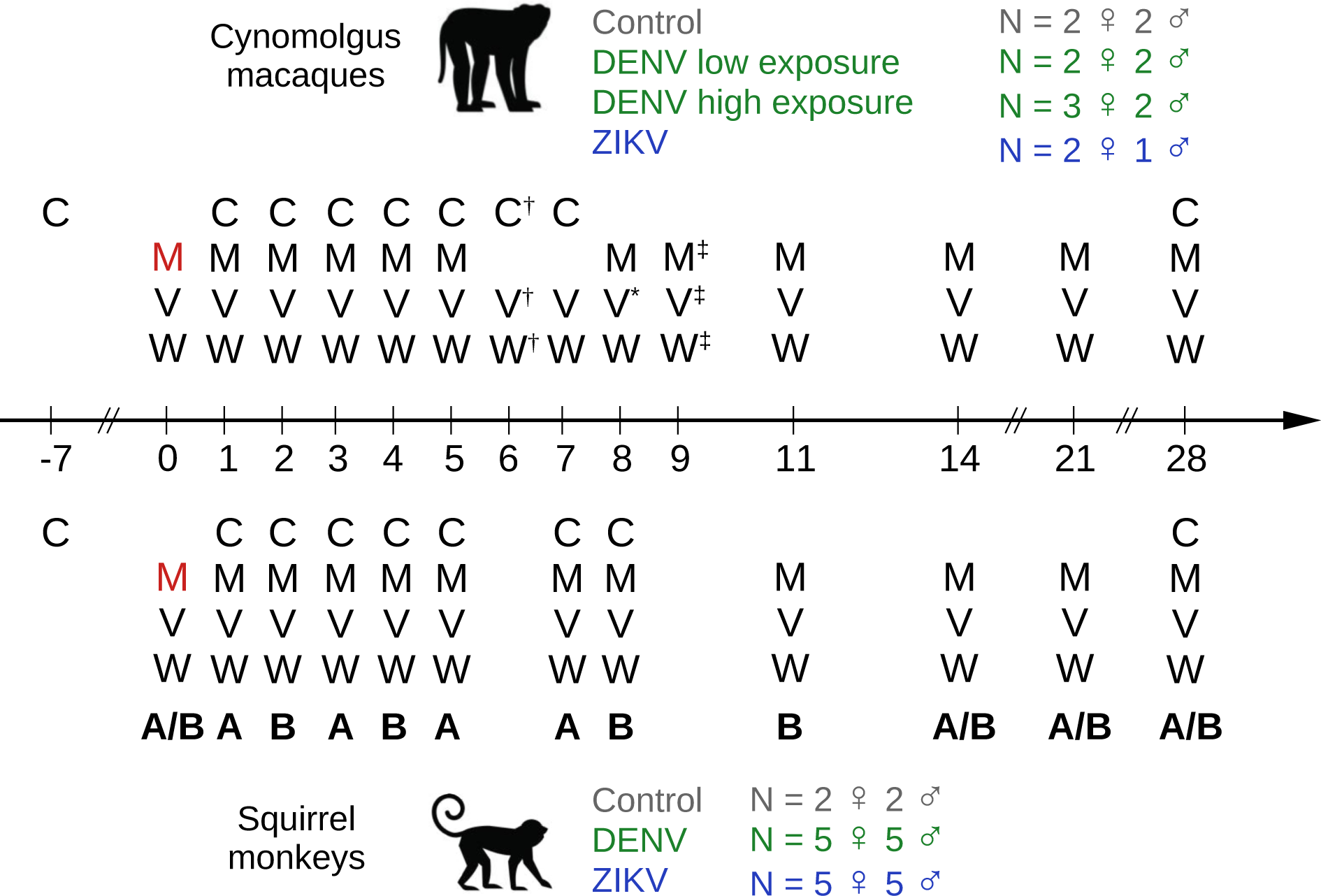
Overview of experimental infections of cynomolgus macaques and squirrel monkeys with sylvatic dengue or Zika virus and subsequent sampling. C: serum sampled for cytokine quantification; M: mosquito feeding (red denotes intrathoracically inoculated mosquitoes used for exposure); V: serum sampled for viremia quantification; W: weight. A and B indicate cohorts, in the case of squirrel monkeys. The monkey images are licensed from Shutterstock. ^†^: only measured in control and DENV-2 infected cynomolgus macaques. ^∗^: for DENV-2 infected cynomolgus macaques, viremia was deduced from transmission to mosquitoes on day 8 and/or viremia on previous and subsequent sampling (see main text). ^‡^: only measured for ZIKV infected cynomolgus macaques.

Adult *Aedes albopictus* mosquitoes were maintained at 28°C while the temperature of the rearing room (from larval development to adult emergence) varied between 26 and 28°C. Temperature for NHP maintenance was 26-28°C for squirrel monkeys and 24-26°C for cynomolgus macaques. All raw data on mosquito engorgement on NHPs are provided in Table S.1.

### Feeding of infected *versus* uninfected mosquitoes: Experiment Day 0

For macaques, on day 0 of the experiment, screen-top, cardboard cartons (diameter 8 cm, height 8.5 cm) containing either 1 (low dose) or 10 (high dose) *Ae. albopictus* (Galveston 2018, F12) that had been intrathoracically inoculated with a sylvatic Malaysian strain of DENV-2 (P8-1407) ten days prior, 15 *Ae. albopictus* intrathoracically inoculated with sylvatic African strain of ZIKV (DakAr 41525) ten days prior, or cartons of 10 uninfected control mosquitoes that had not been intrathoracically inoculated were placed upon the ear of a NHP for an average of 7.5 ± 0.62 minutes (mean ± 1 SE, range = 4 to 10 minutes). For squirrel monkeys, cartons containing either 15 *Ae. albopictus* (Galveston 2018, F14) intrathoracically inoculated with a sylvatic Asian strain of DENV-2 (P8-1407) ten days prior, 15 *Ae. albopictus* intrathoracically inoculated with a sylvatic African strain of ZIKV (DakAr 41525) ten days prior, or 15 uninfected control mosquitoes that had not been intrathoracically inoculated, were placed upon the ear of a NHP for an average of 6.5 ± 0.29 minutes (mean ± 1 SE, range = 4 to 9 minutes). Additionally, to test whether non-viremic transmission might occur by co-feeding, uninfected mosquitoes (n=15) were allowed to feed on squirrel monkeys immediately after they had been fed upon by infected mosquitoes, on the ear not used for the original feeding. These are hereafter termed “co-feeding mosquitoes”. Co-feeding mosquitoes were held upon the ear for an average of 4.9 ± 0.22 minutes (mean ± 1 SE, range = 3 to 6 minutes). Cartons were then returned to an arthropod containment insectary space (ACL2), mosquitoes were cold anesthetized, and visibly engorged mosquitoes were separated, counted, and returned to an incubator, while unengorged mosquitoes were counted and discarded. For the purposes of this study, only visibly engorged mosquitoes are considered to have fed. For analyses, the engorgement rate was defined as the ratio of the number of engorged mosquitoes over the total number of mosquitoes in the carton.

### Statistical analysis of the impact of mosquito infection on engorgement

We tested whether the infection status of mosquitoes affected their engorgement rate on day 0 of the experiments. As none of the co-feeding mosquitoes became infected (Table S.2), we aggregated the cofeeding and control mosquitoes. We excluded mosquitoes belonging to the low dose group (1 mosquito per NHP) feeding on cynomolgus macaques, as they all fed. We also tested differences in engorgement between host species for a given mosquito status (control, DENV-2 or ZIKV infected). Model selection focused on choosing between binomial and betabinomial error distributions (both with a logit link function), the latter accounting for dispersion anomalies, as well as between a generalized linear model (fixed effects only) and a generalized mixed effect model, incorporating monkey ID as a random effect (intercept). Duration of mosquito exposure was used as an offset, *i.e* a scaling factor fixed at 1 reflecting that the tendency to engorge is influenced by the duration of exposure. The models were fitted using maximum likelihood and the selection was done through inspection of models’ residuals, likelihood ratio tests, and corrected Akaike Information Criterion (AICc) comparison. We confirmed that significant results remained as such after using the Benjamini-Hochberg correction for *P* values, with a false discovery rate (false positives / (false positives + true positives)) of 5% [31], but we report the initial *P* values in the results section. When showing predicted probabilities of engorgement from the model, we assume a common duration of 6 minutes, corresponding to the median of all durations.

### Feeding of uninfected mosquitoes on infected *versus* uninfected hosts: Experiment Days 1-28

For both host species, blood was drawn at designated intervals from day 1 to 28 after the initial exposure to mosquitoes to quantify infectious viremia via serial dilution and immunostaining (see [30] for details) and cytokine concentrations (Figure 1). Clarified serum from each animal was subjected to the Cytokine 29-Plex Monkey Panel (ThermoFisher Scientific, Waltham, MA, USA), measuring TGF-β, G-CSF, RANTES, Eotaxin, MIP-1α, GM-CSF, MIP-1β, MCP-1, HGF, VEGF, IFN-γ, MDC, I-TAC, MIF, TNF-α, IP-10, MIG, and interleukins (IL) 1β, 1RA, 2, 4, 6, 8, 10, 12, 15, 17. Each sample was run in duplicate according to manufacturer instructions and average values were ascertained via the standard curve generated for each cytokine per run. Baseline concentrations were measured on day −7. Neutralizing antibody titers were measured as 80% plaque reduction neutralization titers (PRNT80) pre-infection and 28 days post-infection, and weight was measured each time a NHP was handled. Due to their small size, squirrel monkeys were assigned to one of two cohorts, sampled on different days (Figure 1). All NHPs were surgically implanted with DST micro-T temperature loggers (Star-Oddi, Garðabær, Iceland) set to record temperature every 15 minutes. Host body temperature at the time of mosquito feeding was linearly interpolated using available temperature records.

For both species, five- to seven-day-old, uninfected *Ae. albopictus* that had been starved of sucrose for 24 hours were allowed to feed on NHP ears (in general n=10 mosquitoes for cynomolgus macaques, n=15 for squirrel monkeys, see Table S.1 for exceptions) on the days designated in Figure 1, and the time of day and duration of exposure were monitored. In two cases, the end time of exposure was not recorded, and instead we used the time the NHP was returned to its cage, which is a slight overestimation. We alternated ears each feeding.

Transmission to mosquitoes was assessed through the presence of virus in either mosquitoes’ bodies or legs or both ([30], Text S.1). This information was sometimes used to correct our measure of infectious viremia (Text S.1).

Squirrel monkeys were euthanized at the termination of this experiment but cynomolgus macaques were not. On day 28 of the experiment, squirrel monkeys were being prepared for euthanasia, which necessitated some changes in the way they were handled, and induced important differences with cynomolgus macaques with regards to time of day and duration of mosquito exposure, which could bias our analyses (Text S.1). We therefore excluded data on day 28 of the experiment, for both species (from n = 331 to n = 293). In this final dataset, mosquito exposure events happened between 8:37 AM and 2:32 PM. Lights were on from 7AM to 7PM to minimize the impact of diel fluctuations on host body temperature. For analyses, the engorgement rate was again defined as the ratio of the number of engorged mosquitoes over the total number of mosquitoes in the carton.

### Statistical analysis of the impact of host infection and biology, as well as time of day, on mosquito engorgement

We tested whether the infection status of NHPs affected uninfected mosquitoes’ engorgement rates. We started with a simple model comparing the experimental groups (combination of NHP species and infection status, and virus species) to see what could be concluded in the absence of other information. Then, to account for heterogeneity within these groups, we built a complete model, accounting for virus species, time of day, host sex, species, body temperature at the time of feeding, weight, and viremia. All quantitative variables were scaled except viremia. Duration of mosquito exposure was used as an offset in both the simple and complete models. The model selection procedure was similar to the one described earlier, except that generalized mixed effect models also included days post infection as random intercepts. Preliminary tests resulted in the inclusion of an interaction term between host species and weight, as well as between host species and host body temperature (Text S.1, Figure S.1). When showing predicted probabilities of engorgement from the model, we assume a common duration of 6 minutes, corresponding to the median of all durations.

### Comparison with inter-individual variation in approach rate of free-living *Aedes albopictus*

In order to compare the inter-individual heterogeneity in our experimental dataset to heterogeneity in the approach rate of free-living mosquitoes, we took advantage of an experiment designed to test the impact of levels of urbanization on the approach of *Aedes* spp. mosquitoes to humans [32]. In this experiment, human approach rate was measured by handheld net collections of mosquitoes that would approach within an arm’s length to the collector. Collectors were wearing mosquito repellent to prevent bites.

To compare experimental and field data, we used the coefficient of variation (standard deviation/mean), which is unitless. We computed it on engorgement rates normalized by duration of exposure, per NHP, for our dataset, and on number of approached females *Ae. albopictus*, per collector (the duration of collection being constant), in what we defined as optimal conditions (Text S.2), for approach data.

### Effect of repeated exposure to uninfected mosquitoes on cytokine response

Lastly, we took advantage of the experimental design of Hanley *et al.* [30] to determine whether the immune response of NHPs was notably affected by the exposure to uninfected mosquitoes. To do so, we assessed the effect of repeated exposure to uninfected mosquitoes on control NHP cytokine concentration (N = 4 control cynomolgus macaques and 4 control squirrel monkeys). We used log_10_ cytokine concentration as the response variable, and variables measuring mosquito exposure as fixed effects (Table S.3). For simplicity, here we use the term bite to refer to the number of engorged mosquitoes at the end of the exposure period, although we did not observe the number of biting events *per se*. We considered the effects of biting in the short term (the number of bites received the day prior for cynomolgus macaques, and 2 days prior for squirrel monkeys, due to alternating days of exposure) and the long term (the cumulative number of bites received in the last seven days for both species). The model selection procedure is detailed in Text S.3.

## Results

Before diving into analyses, we wish to highlight that although mosquitoes were starved from sucrose and blood meal and given no choice of hosts, the engorgement rates ranged from 0% to 100%, including on viremic NHPs (Figure S.2A).

### Impact of mosquito infection status on engorgement rates

In general, mosquito infection status had no significant effect on engorgement rate (Figure 2, Table S.4), save that DENV-2 infected *Ae. albopictus* were less likely to engorge than control *Ae. albopictus* when feeding on squirrel monkeys (odds ratio OR = 0.31 [0.13; 0.71], p = 2.0e-4). We also detected an interaction between host species and mosquito infection status on feeding behavior, in that we found significantly higher engorgement of DENV-2 infected mosquitoes on cynomolgus macaques than on squirrel monkeys (OR = 6.46 [1.65; 25.3], p = 3.2e-4) but no difference between host species for ZIKV infected or control mosquitoes (Table S.4). The selected model was a logistic regression model with a betabinomial error distribution, without random effects (Tables S.5, S.6).

**Figure 2:**
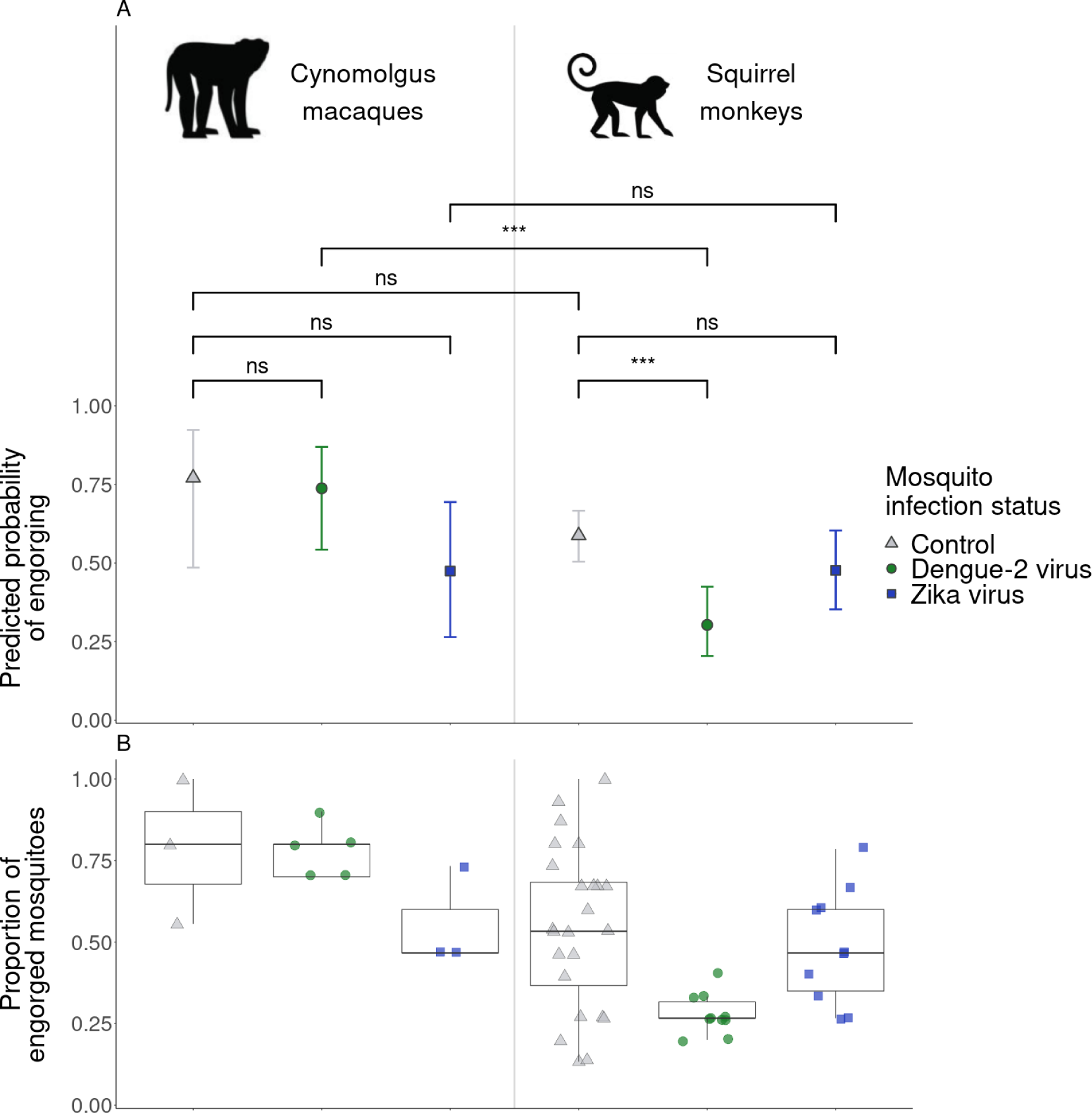
Engorgement rates of *Aedes albopictus* on day 0 of the experiments, depending on the mosquito infection status and NHP species. **A** – Probabilities of feeding predicted by the model per group, assuming a duration of mosquito exposure of 6 minutes (median of all exposures), with results of statistical comparisons indicated above brackets, with ns: p > 0.05, ***: 0.0001 < p < 0.001. **B** – Raw data (one point = one NHP, one day) colored by NHP infection status (grey triangle: control, green circle: DENV-2 infected, blue square: ZIKV infected), with one boxplot per group. Duration of mosquito exposure varies but is not shown here. Data from day 28 excluded (see Methods).The monkey images are licensed from Shutterstock.

### Impact of host infection status and host species on engorgement rate

In a simple model, NHP infection status had generally no significant effect on engorgement rate (Figure 3A,B, Table S.7), save that *Ae. albopictus* were less likely to engorge on ZIKV infected cynomolgus macaques than on control cynomolgus macaques (OR = 0.44 [0.23; 0.85], p = 7.8e-4). When restricting the dataset to the range of days where viremia was detected for each virus (days 1-14 for DENV-2, 1-8 for ZIKV), the results remained qualitatively unchanged (Figure 3C). The selected simple model used a betabinomial error distribution without random effects (Tables S.8, S.9).

**Figure 3:**
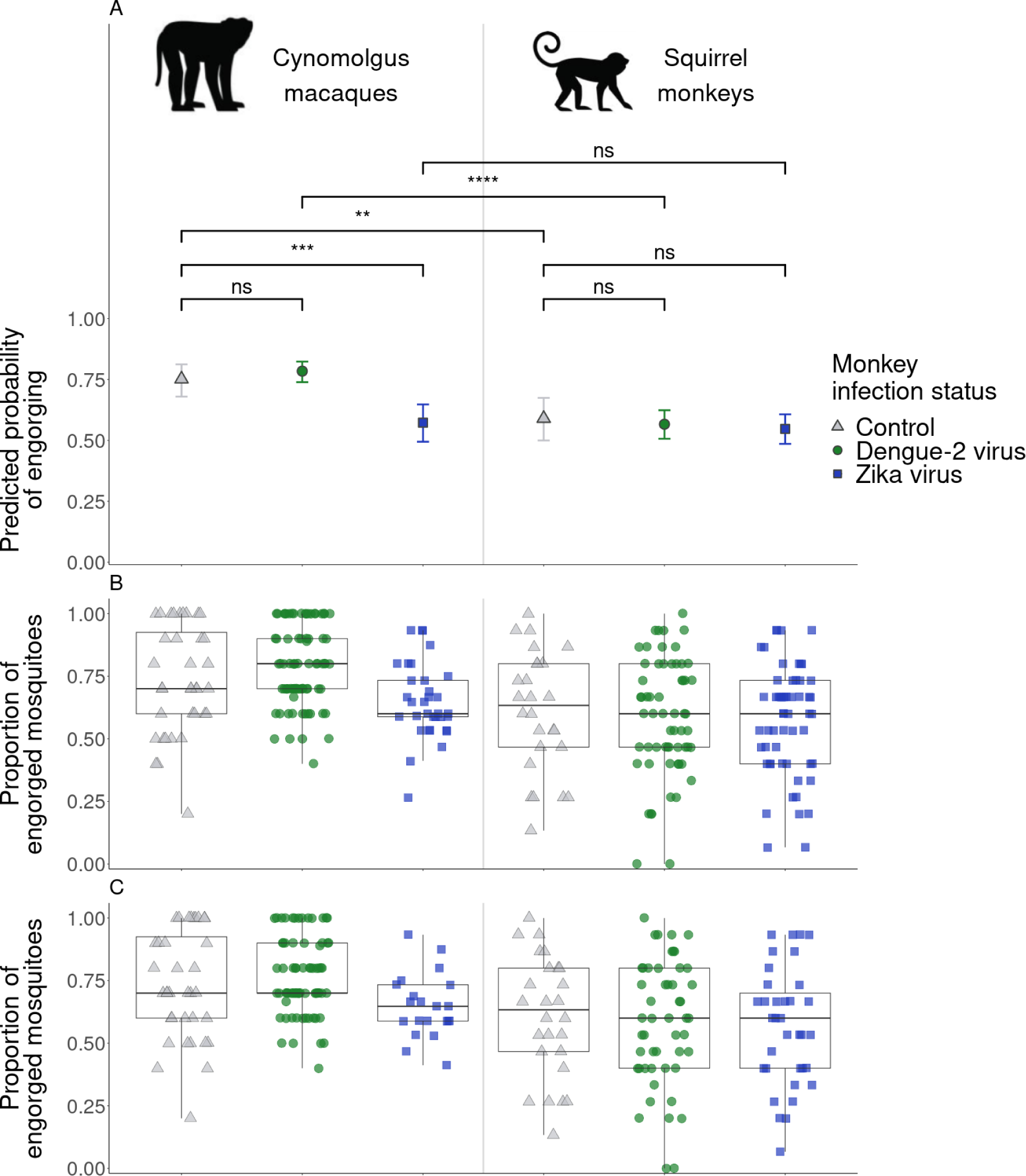
Engorgement rates of uninfected *Aedes albopictus* depending on NHP infection status and species. **A** – Probabilities of feeding predicted by the simple model per group, assuming a duration of mosquito exposure of 6 minutes (median of all exposures), with results of statistical comparisons indicated above brackets, with ns: p > 0.05, **: 0.001 < p < 0.01, ***: 0.0001 < p < 0.001, ****: p < 0.0001. **B** – Raw data (one point = one NHP, one day) colored by NHP infection status (grey triangle: control, green circle: DENV-2 infected, blue square: ZIKV infected), with one boxplot per group. Duration of mosquito exposure varies but is not shown here. Data from day 28 excluded (see Methods). **C** - Raw data, same as B except that for infected groups, we restrict the data to the range of days where viremia was detected for each virus (days 1-14 for DENV-2, 1-8 for ZIKV). The monkey images are licensed from Shutterstock.

Furthermore, we found significantly higher *Ae. albopictus* engorgement rates on cynomolgus macaques than squirrel monkeys when considering control NHPs (OR = 2.10 [1.05; 4.22], p=0.0043) or DENV-2 infected NHPs (OR = 2.78 [1.74; 4.45], p = 5.8e-9), but not ZIKV infected NHPs (p = 0.61).

### Impact of host infection and biology as well as time of day on engorgement rate

The complete model tested the effect of host species, sex, weight, body temperature at the time of feeding, virus and viremia, as well as time of day, on engorgement rates. Two-way interactions between host species and weight, and host species and body temperature, were included. The model highlighted a significant, positive effect of host body temperature on *Ae. albopictus* engorgement on squirrel monkeys (p = 0.019), but not on cynomolgus macaques (p = 0.90). The relationship between host body temperature and engorgement rates is showed in Figure 4. No other variables had a significant effect (Table S.10). The selected complete model used a betabinomial error distribution and included random effects for monkey ID and day (Tables S.11, S.12).

**Figure 4:**
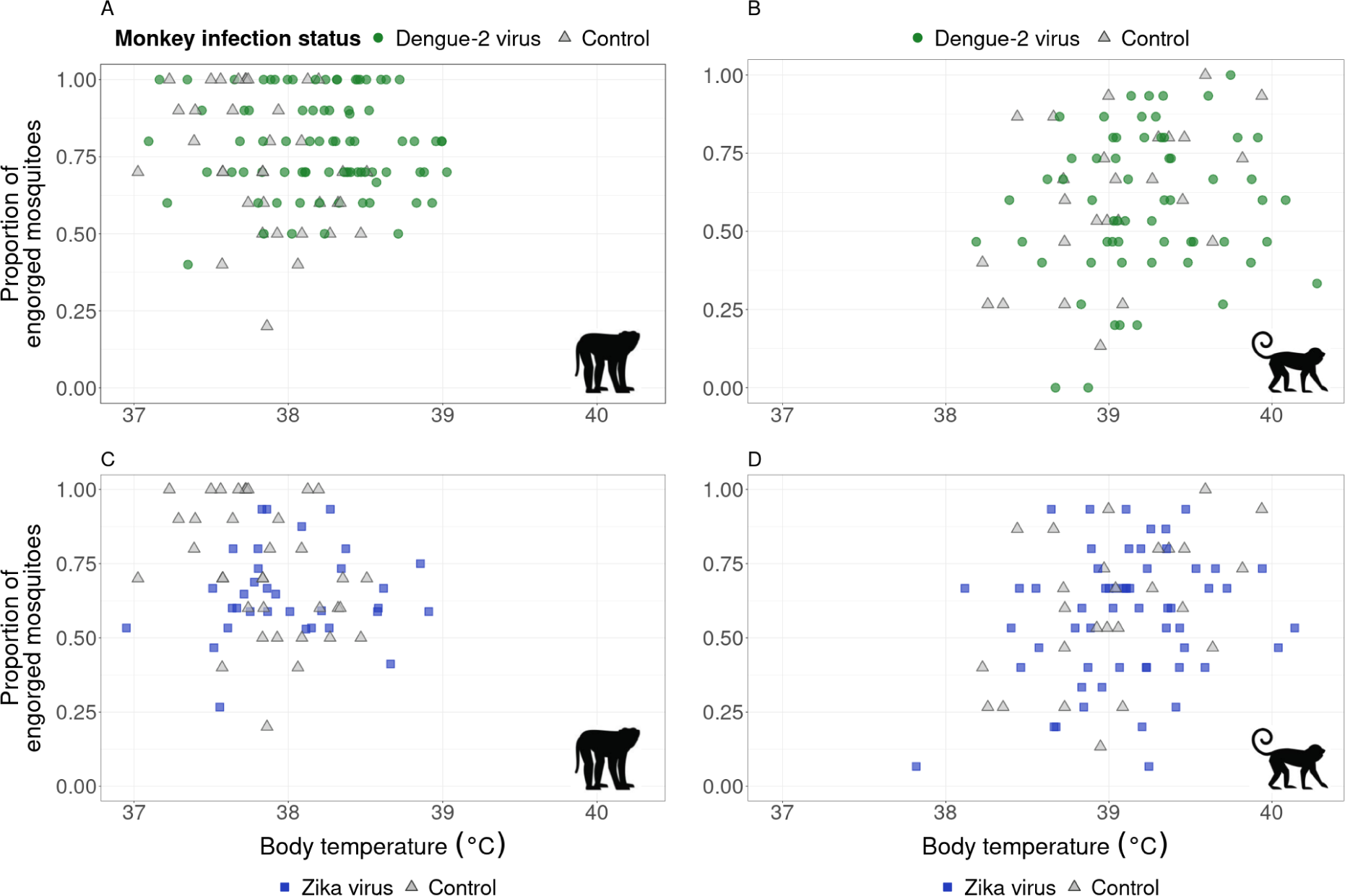
Relationship between host body temperature and *Aedes albopictus* engorgement rates, in cynomolgus macaques (left column) and squirrel monkeys (right column). Raw data (one point = one NHP, one day) colored by NHP infection status (grey triangle: control, green circle: DENV-2 infected, blue square: ZIKV infected). Note that the same set of control animals are shown for same species experiments (bottom and right panels of a given column). The duration of mosquito exposure varies but is not shown here. Data from day 28 excluded (see Methods). The monkey images are licensed from Shutterstock.

### Degree of inter-individual variation in mosquito engorgement rates among hosts in this study compared to approach rates in free-living *Aedes albopictus*

In our experiments, engorgement rates varied substantially within and between NHPs (Figure 5). The coefficient of variation per NHP ranged from 0.22 to 0.88, with a mean of 0.45. In the study measuring the approach of free-living mosquitoes to humans [32], the number of approaching female *Ae. Albopictus* ranged from 0 to 6 (Figure S.2B) and the coefficient of variation ranged from 0 to 2, with a mean of 1.43.

**Figure 5:**
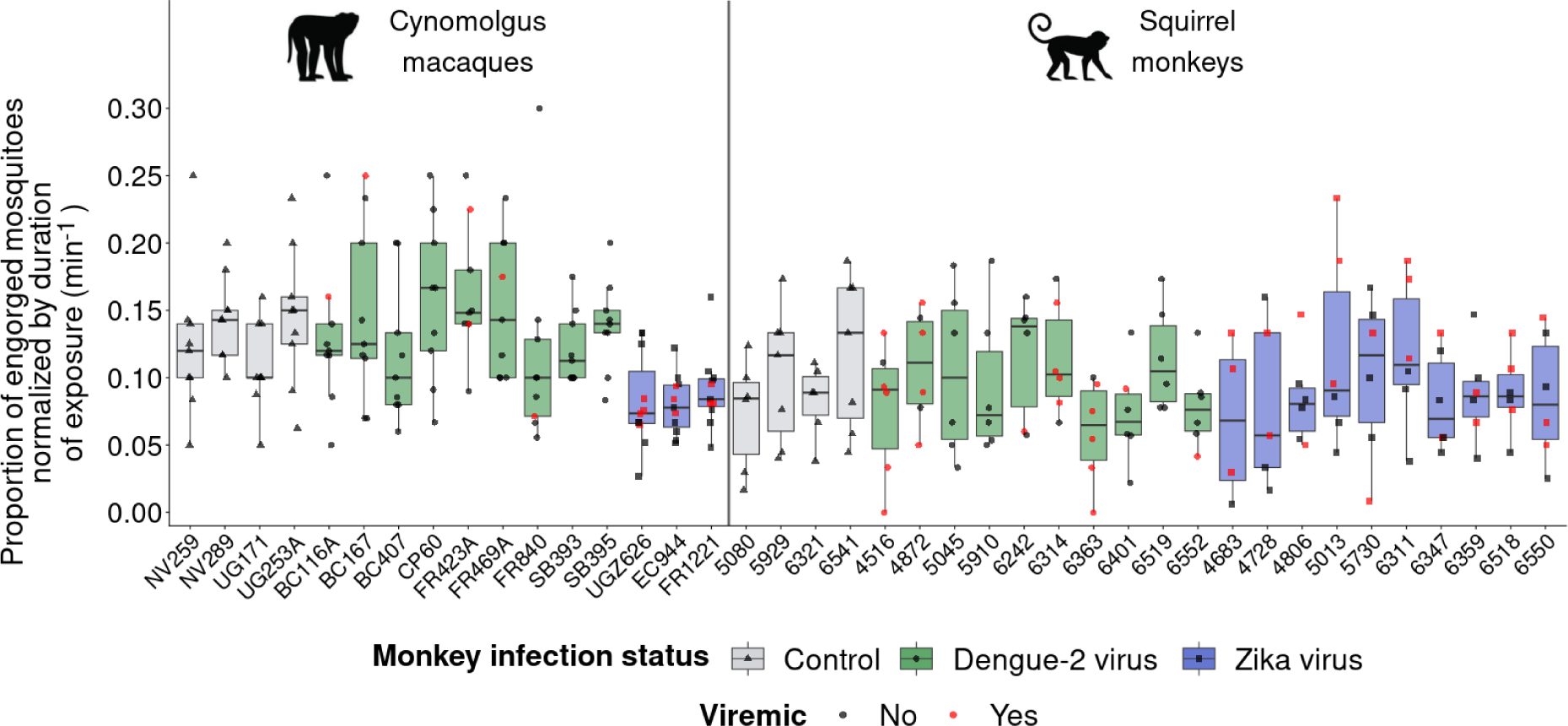
Engorgement rates of *Aedes albopictus* on individual NHPs, normalized by duration of exposure, depending on monkey infection status and species. One boxplot per NHP, colored by NHP infection status (grey: control, green: DENV-2 infected, blue: ZIKV infected), with raw data (triangle: control, circle: DENV-2 infected, square: ZIKV infected). Red points indicate feedings on animals that were detectably viremic, or that resulted in transmission to mosquitoes. Data from day 28 excluded (see Methods). The monkey images are licensed from Shutterstock.

### Repeated exposures to bites of uninfected mosquitoes shaped EGF and MIF concentrations in cynomolgus macaques

We considered the effects of mosquito bites on cytokine concentrations in control NHPs, both in the short term (the number of bites received the day prior for cynomolgus macaques, and 2 days prior for squirrel monkeys, due to alternating days of exposure) and the long term (the cumulative number of bites received in the last seven days for both species). We included all cytokine measures from day −7 to day 28, and varied the short-term and long-term exposure variables accordingly, accounting for days with no mosquito bites.

In control cynomolgus macaques, daily epidermal growth factor (EGF) and macrophage migration inhibitory factor (MIF) concentrations were significantly associated with short-term and long-term exposure to uninfected mosquito bites. For EGF, the concentrations increased by 0.02 [0.01; 0.03] log_10_ pg/µl per additional bite the day prior (p = 2.15e-4, Figure 6A, Text S.3, Table S.13). For MIF, the relationship with short-term exposure was non-linearly positive, with a saturation after about 6 bites the day prior (p = 0.005, Figure 6B, Text S.3, Table S.13). Both cytokines decreased linearly with long-term bite exposure (−0.006 [−0.008; −0.004] log_10_ pg/µl per additional bite in the last seven days for EGF, p = 8e-6, −0.010 [−0.014; −0.005] for MIF, p = 2.3e-4, Figure 6C,D, Text S.3, Table S.13). Daily tumor growth factor β (TGF-β) and monocyte chemoattractant protein-1 (MCP-1, also called CCL2) concentrations were also significantly, negatively associated with long-term bite exposure (Text S.3, Figure S.3, Table S.13). All measures were above the limit of detection for these cytokines.

**Figure 6:**
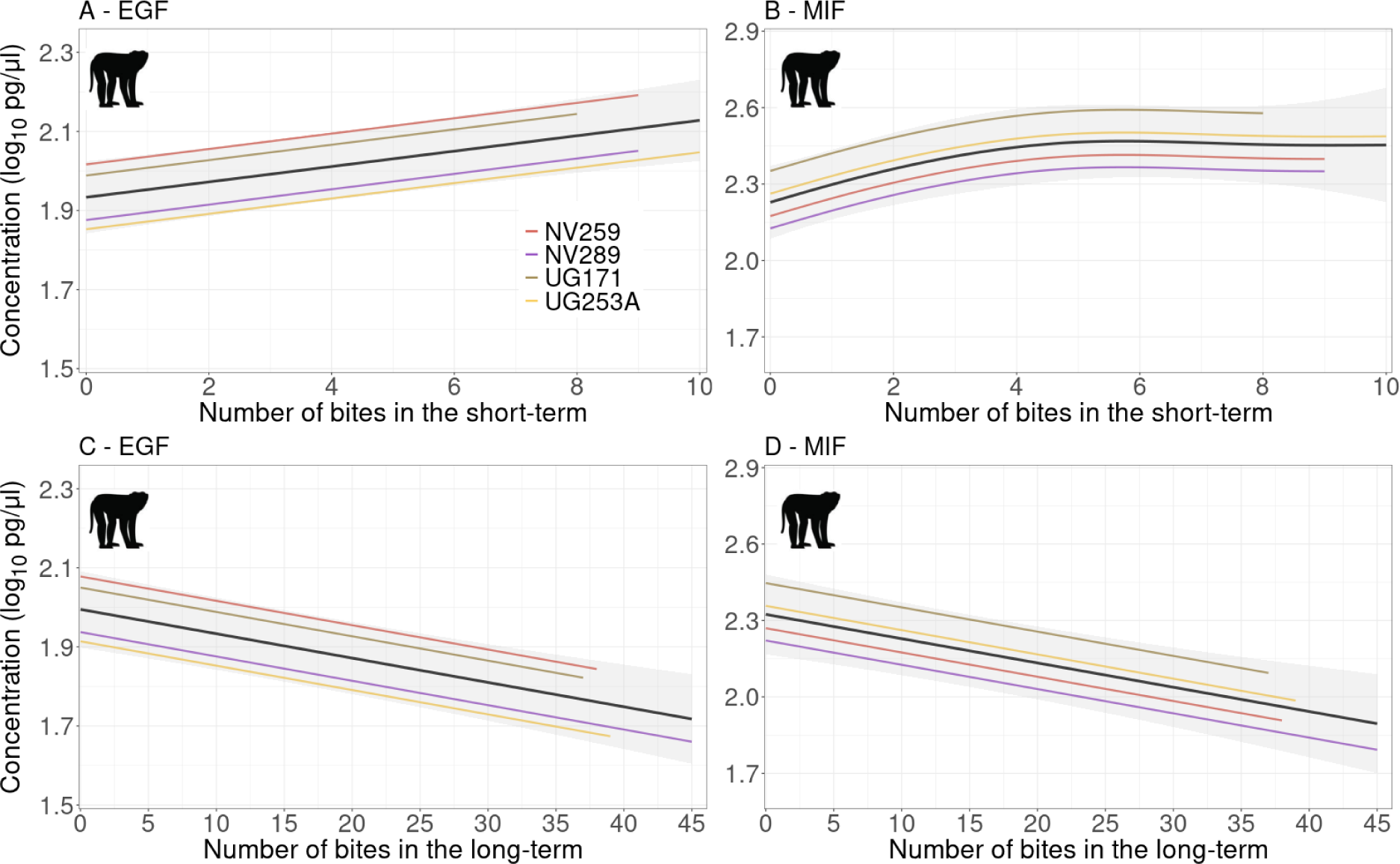
In control cynomolgus macaques, concentrations of cytokines EGF (A,C) and MIF (B,D) are significantly associated with the number of uninfected mosquitoes which engorged the day before (A,B) and in the last 7 days (C,D). Results from generalized additive mixed effect models, estimating an intercept per NHP. Thick black lines and shading show the population trends with uncertainty. Colored lines are individual fits. Fits for a given exposure variable (short-term or long-term) are computed with the other variable fixed (0 for short-term, 10 for long-term), which is why datapoints are not plotted (short-term and long-term variables varied concomitantly in the experiments). Note that y-axis scales differ between cytokines (A,C *vs.* B,D), and that x-axis scales differ between exposure variables (A,B *vs.* C,D). The monkey images are licensed from Shutterstock.

In cynomolgus macaques, interleukins 1β, 4, 5, 10, 17, and VEGF could not be tested because all measures were below the limit of detection. All other tested cytokines showed no significant association with bite exposure.

In squirrel monkeys, we could only test I-TAC, MIF, MIP-1β and RANTES, as all other cytokines had too many observations below the limit of detection. These 4 cytokines showed no significant association with bite exposure.

## Discussion

Identifying the factors that shape host choice by arbovirus vectors is critical for advancing model-based predictions of arbovirus spillover, spillback, and among-host transmission. Here, we studied the engorgement rates of *Ae. albopictus*, a likely bridge vector between sylvatic and urban cycles of arbovirus transmission, on two non-human primate species, investigating the effect of both vector and host infection with sylvatic strains of DENV-2 or ZIKV.

We did not find any systematic effect of vector infection with DENV-2 or ZIKV on the feeding behavior of *Ae. albopictus*, save for a tendency of DENV-2 infected *Ae. albopictus* to engorge at lower frequency on squirrel monkeys than control *Ae. albopictus*. Wei Xiang *et al.* [20] showed that DENV-2 infected *Ae. aegypti* were more attracted to uninfected mice than control mosquitoes, but were less successful when probing, necessitating more probes to feed to repletion. In combination, these two effects resulted in similar blood-feeding rates between infected and control mosquitoes, over an observation period of 30 minutes. Because mosquitoes in our experiments were only given a limited time to engorge (due to the need to minimize time under anaesthesia for the NHPs), impaired probing could lead to lower rates of engorgement, even if infected mosquitoes were more attracted to hosts. We note that we were not able to observe probing in our experiments. Additionally, in the Wei Xiang *et al.* study, mosquitoes were infected by feeding on a bloodmeal containing virus, recapitulating all steps of mosquito infection, whereas in our study, mosquitoes were infected by intrathoracic inoculation, a procedure that bypasses the mosquito midgut. Furthermore, in our experiments, control mosquitoes were not subjected to intrathoracic injection, which could have confounded effects of infection on feeding. However, intrathoracic inoculation was conducted 10 days prior to feeding on NHPs, giving mosquitoes substantial time to recover from the procedure.

We also did not find any universal effect of host infection with sylvatic DENV-2 or ZIKV on *Ae. albopictus* engorgement rates, with the exception that mosquitoes engorged significantly less frequently on ZIKV infected macaques than control macaques. These results contrast with those of a recent study by Zhang *et al.* [14]. In their study, Zhang *et al.* first used a dual-choice olfactometer assay to establish that *Ae. aegypti* and *Ae. albopictus* were more attracted to host cues produced by DENV-2 or ZIKV infected mice or humans relative to controls, but mosquitoes were not allowed to feed. These results cannot be compared to ours because of differences in protocol. However in another experiment, *Ae. aegypti* were allowed to directly access mice, and this experiment also found increased attraction to infected hosts. Differences in the findings of Zhang *et al.* and the current study may reflect differences in the vector species used (*Ae. aegypti* in [14], *Ae. albopictus* in the present study), the host species used (immunodeficient mice in [14], natural NHP hosts in the present study), the ecotype of virus used (human-endemic in [14], sylvatic in the present study), or host infection route (needle delivery in [14], mosquito bite in the present study). Cozzarolo *et al.* [13] suggested that the ideal method for interrogating mosquito feeding preferences would be dual-choice experiments where vectors are allowed to bite, and hosts allowed to defend themselves, but this is not possible when protocols require animals to be anesthetized, as was the case here and in Zhang *et al.*. Nevertheless, in our experiments NHPs did not display any signs or behavior associated with illness [30], which could have driven a possible preference towards infected animals. Moreover, Buchta *et al.* [33] showed that anaesthesia, by dropping core body temperature, could impact mosquito feeding, in experiments performed on guinea pigs with *Anopheles stephensi* and *Phlebotomus papatasi*. They recommended the use of a warming device to maintain normothermic body temperature of the host when conducting feeding experiments with mosquitoes. In our case, core body temperature was maintained in squirrel monkeys by placing them on a preheated warming blanket. This was not done for cynomolgus macaques, but mosquito exposure occurred shortly after the anaesthesia so that their temperature had not yet dropped significantly. Importantly, the anesthesia procedure for control and infected monkeys was identical, so while anesthesia could have affected overall feeding rate it could not have biased comparisons. It will be important to disentangle the possible long-range and short-range effects of host infection status on mosquito attraction and actual bites, possibly with a similar framework to Wei Xiang *et al.* [20], to see how these effects impact the overall contact rate and opportunities for virus transmission.

A recurrent effect on mosquito engorgement in our study was host species, with a preference towards cynomolgus macaques, which was evident in control NHPs and DENV-2 infected NHPs but not in ZIKV infected NHPs. In day 0 mosquitoes, this species difference was only seen from DENV-2 infected mosquitoes, but this could be due to smaller sample sizes. In natural settings, *Ae. albopictus* is known to feed on a wide range of hosts [34], and preferences can be driven by visual cues such as body size [35]. However, in our set-up mosquitoes could not detect hosts from a distance, and weight (a proxy of body size) did not explain observed engorgement rates according to our model. To interpret differences in attractiveness between NHP species, other host-level factors such as host odor [8, 9], which is in part shaped by skin microbiome [36], should be investigated. Whether host or vector infection can counter-balance a possible trophic preference of *Ae. albopictus* towards certain NHP species needs to be investigated further.

After accounting for other possible drivers of engorgement, the only variable that stood out was host body temperature, with a positive effect in squirrel monkeys only. Studies usually focus on mosquitoes’ ability to detect warm-blooded hosts from a background temperature [37, 38], rather than the effect of host body temperature itself. Cynomolgus macaques’ body temperature was on average 1.2°C lower than squirrel monkeys’ at the time of mosquito feeding, but cynomolgus macaques were housed in rooms on average 2°C lower than squirrel monkeys. Higher body temperature has been suggested as a possible explanation of why pregnant women were more attractive to malaria vectors than non-pregnant women [39], with an average difference of temperature between their abdomens of 0.7°C. In the present study, squirrel monkeys’ body temperature ranged from 37.82°C to 40.28°C at the time of mosquito feeding, and the associated predicted probabilities of engorging ranged from 0.48 to 0.76. A positive effect of body heat on mosquito engorgement might be linked to an increased release of volatile compounds [39]. However, in our case, we emphasize that the effect of host body temperature was weakly supported. Rather, our study depicts mosquito engorgement as a highly variable, multi-factorial phenomenon [40], even in controlled conditions.

In a context of disease transmission, focusing on a single event per mosquito batch, namely the proportion of engorged mosquitoes, as we did, following a relatively long exposure to a host, might not be sufficient. This outcome has the advantage of being easy to measure but does not represent the feeding behavior in its entirety. Other metrics such as the time to first bite, time between bites, duration of probing and feeding, have been showed to play a role in pathogen transmission rate [15, 20, 41]. Several mathematical models have sought to finely represent the biting behavior of mosquitoes and its influence on transmission dynamics [41–44]. Nonetheless, in epidemiological models of vector-borne diseases transmission dynamics, the biting rate is most often represented as a constant, neither influenced by abiotic (e.g temperature) nor by biotic (e.g host species, infection status, or allometry, [45, 46]) factors. Another common modelling hypothesis is to assume that a unique blood meal takes place per gonotrophic cycle, whereas imbibing multiple blood meals is a common behaviour in multiple species, including *Ae. aegypti* [47, 48]. This has important implications, as recent studies have showed that successive feeding episodes could enhance viral dissemination within mosquitoes [49–51]. Vector preference strategies, particularly in multi-host systems, can have important epidemiological consequences, which can be studied conceptually through mathematical models [52, 53]. More studies are needed, both in controlled experimental settings and in the field, to understand the underlying factors that influence mosquito biting habits.

The variability of *Ae. albopictus* engorgement per NHP and between NHPs was striking in our experiments, and the sometimes-low engorgement rates observed were surprising to us as mosquitoes had no other choice of host. This variability was less extreme compared to the variability among collectors when recording approach from free-living *Ae. albopictus* in the field. Still, both controlled and field experiments point towards the importance of incorporating individual-level heterogeneity of biting in models at the population scale, even in the absence of mechanistic drivers for now.

When they bite, mosquitoes deliver salivary gland proteins, a complex cocktail that can impact host immunity [54] and, when a virus is also delivered, its viral dynamics [55–57]. While the impact of mosquito saliva on host susceptibility to arboviruses and subsequent arbovirus dynamics has been well studied [58], particularly in mice, the cytokine response of NHPs to the bite of uninfected mosquitoes has received relatively little attention. We took advantage of data collected in the current study to begin to close this knowledge gap. In control cynomolgus macaques, among 29 cytokines analyzed, we identified short-term positive associations between uninfected mosquito bites and EGF and MIF concentrations, as well as long-term negative associations with EGF, MIF, TGF-β, and MCP-1 concentrations, while the remaining 25 cytokines assayed showed no significant responses to repeated mosquito bites. This is consistent with recruitment of leukocytes to the bite site (EGF, MCP-1) to resolve inflammation (MIF, TGF-β), which slows down over time as bites keep happening, but the response is already in place. In their study on humanized mice bitten by uninfected mosquitoes, Vogt *et al.* [54] noticed a decrease in MCP-1 concentrations 6 hours post-bite and an increase 7 days post-bite. They observed no change in EGF 24 hours post-bite, and did not measure MIF and TGF-β. Here, we focused on the cytokine response in the absence of pathogen transmission, but in studies of infected animals, IL-4 and IL-10 are often cited as being upregulated in response to arbovirus infection through mosquito bites [55–57, 59]. In our study, control NHPs of both species produced concentrations of these two cytokines below the limit of detection over the whole duration of the experiment, suggesting that this upregulation does not happen in the absence of arbovirus transmission. Although cynomolgus macaques were housed indoors for several months before the start of the study, they initially came from Mauritius and therefore likely had previous exposure to *Ae. albopictus*. Future experiments could pursue skin biopsies to investigate the immediate mobilization of these cytokines following a mosquito bite [60].

## Data and code accessibility

All analyses were performed in R. Data, scripts, and outputs are available at https://github.com/helenececilia/albopictus_engorgement_DENV_ZIKV_monkeys.

## Acknowledgements

This research was funded by grant 1R01AI145918-02 (to KAH, NV, BMA) and in part by the Centers for Research in Emerging Infectious Diseases “The Coordinating Research on Emerging Arboviral Threats Encompassing the Neotropics (CREATE-NEO)” grant U01AI151807 from the U.S. National Institutes of Health (to N.V. and K.A.H). Squirrel monkeys acquired from the UT MD Anderson Cancer Center, Michael E. Keeling Center for Comparative Medicine and Research were supported by NIH grant 5P40OD010938-36 prior to sale to UTMB. Fellowship 709139 from CONACYT (Consejo Nacional de Ciencia y Tecnología, Government of Mexico) was used to produce data on mosquito approach in the field in Manaus, Brazil.

## Author contributions

**Conceptualization**: H.C, B.M.A, N.V, K.H.A. **Methodology**: H.C, B.M.A, K.H.A. **Investigation**: S.R.A, B.M, R.Y, K.H.A, N.V., S.L.R. **Visualization**: H.C. **Formal analysis**: H.C. **Software**: H.C. **Funding acquisition**: N.V, K.H.A. **Project administration**: N.V, K.H.A., S.L.R **Supervision**: B.M.A, K.H.A. **Writing original draft**: H.C, B.M.A, K.H.A. **Writing review and editing**: H.C, B.M.A, S.R.A, B.M, R.Y, S.L.R, N.V, K.H.A. All authors contributed to the article and approved the submitted version.

