## Supplementary Information for "*Aedes albopictus* is not an arbovirus aficionado – Impacts of sylvatic flavivirus infection in vectors and hosts on mosquito engorgement on non-human primates"

Note : Tables S.1, S.2, S.3, and S.13 are provided as separate files.

#### S.1 Preliminary observations

For each carton, engorged and unengorged mosquitoes were counted and separated; engorged mosquitoes were returned to an incubator for 14 days and then bodies and legs were separated and assayed for virus infection as described above. Transmission to mosquitoes was assessed through the presence of virus in either mosquitoes' bodies or legs or both (Hanley *et al.* 2023). In some cases, NHPs with no detectable viremia, even after sera were passaged in Vero cells, nonetheless transmitted virus to mosquitoes. In these cases, we assigned a viremia of 10 PFU/ml (half the limit of detection) to the monkey. Similarly, if a NHP was not detectably viremic nor transmitted to mosquitoes on a given sampling day, but was viremic and/or transmitted to mosquitoes on the previous and following sampling days, we also assigned a viremia of 10 PFU/ml. The remaining samples with no detectable viremia even after passage, and no transmission to mosquitoes were considered as true absence of viremia. A cynomolgus macaque transmitted virus to mosquitoes on day 8 and was assigned a viremia of 10 PFU/ml, in the absence of actual viremia measurement (due to a staggered sampling of serum and mosquito feeding in this one case).

Squirrel monkeys were euthanized at the termination of this experiment but cynomolgus macaques were not. On day 28 of the experiment, squirrel monkeys were being prepared for euthanasia, which necessitated some changes in the way they were handled. Thus, on day 28, mosquito exposure happened significantly later than on other days in squirrel monkeys ($p < 2e\text{-}16$), but not in cynomolgus macaques ($p = 0.12$). On day 28, exposure also lasted significantly longer than on other days in squirrel monkeys (8 to 15 minutes, with a median of 11 minutes for day 28, and 4 to 12 minutes, with a median of 6 minutes for other days, $p = 1.7e\text{-}9$), but not in cynomolgus macaques (5 to 8 minutes, with a median of 6 minutes for day 28, and 3 to 16 minutes, with a median of 6 minutes for other days, $p = 0.77$). To avoid skewing the distribution of these time metrics, our models were fitted without data from day 28 for both species (from $n = 331$ to $n = 293$).

The following tests were performed to decide on the inclusion of interaction terms in our complete model. Data for days 0 and 28 were excluded. Tests on temperature were performed using a generalized linear mixed effect model with monkey ID and day as random intercepts. Tests on weights used a linear model, and tests on duration of exposure used a generalized linear model with a Poisson error distribution.

Over the whole course of the experiments, temperatures were lower in cynomolgus macaques than in squirrel monkeys (Figure S.1), both in control ($p = 0.002$), DENV-2 infected NHPs ($p = 0.0004$), and ZIKV-infected animals ($p = 0.001$). This species difference was still apparent when restricting to temperatures at the time of mosquito exposure, and in addition, temperatures were higher in DENV-2 infected than control cynomolgus macaques (Figure S.1, $p = 0.022$). The time at which mosquito exposure happened influenced host temperature (Figure S.1). In cynomolgus macaques, we notice signs of hypothermia (Figure S.1A,B), likely due to anesthesia, and because of the absence of heating blanket, but mosquito exposure mostly happened before such hypothermia (Figure S.1A,B). We also note that within monkey species, the variability of temperature at the time of mosquito exposure was limited to a range of about 2°C (Figure S.1). We included an interaction term between temperature and species in our complete model of mosquito engorgement on days 1-21.

Squirrel monkeys were lighter than cynomolgus macaques ($p < 2$ e-16, see Hanley *et al.* 2023). We therefore included an interaction term between weight and species in our complete model of mosquito engorgement on days 1-21.

The duration of mosquito exposure lasted on average longer on ZIKV-infected macaques than control cynomolgus macaques (6 minutes [5 ; 7] for controls, 8 minutes [7 ; 9] for ZIKV-infected, $p = 0.0008$).

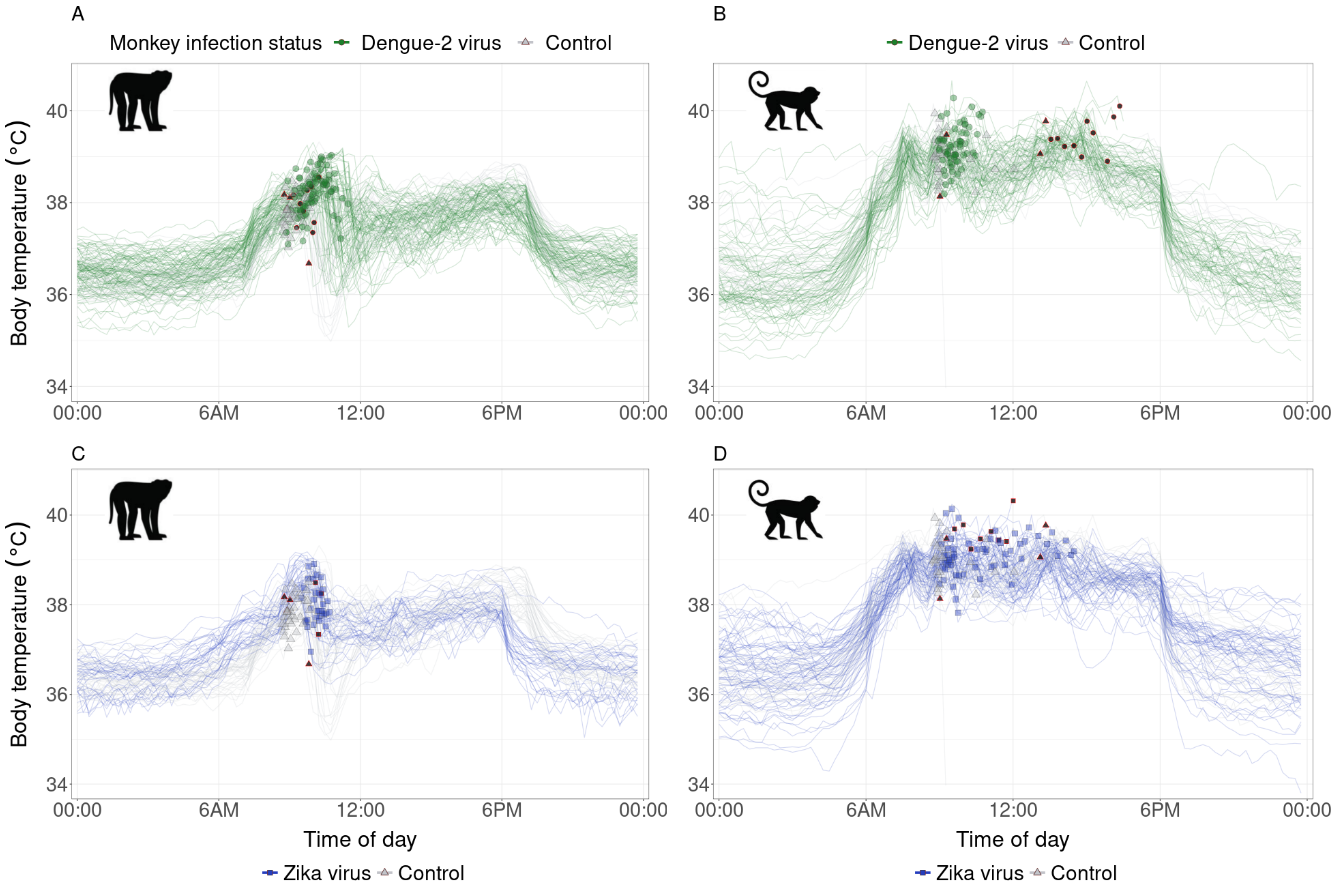


Figure S.1: Body temperature dynamics of monkeys on days of mosquito exposure (lines), recorded through transponders, with a focus on temperature at the time of mosquito exposure (points). Day 0 of the experiments is excluded here, as well as temperatures after the last exposure on day 28. **A, C** : Cynomolgus macaques. **B, D** : Squirrel monkeys. Green denotes DENV-2 infection, blue ZIKV-infection, and grey is for control monkeys. Note that the same set of control animals are shown for same species experiments (left and right columns). Smaller black points denote day 28 data, which was excluded from analyses (see main text).

### Feeding of infected and uninfected mosquitoes : Experiment Day 0

| Comparison | Estimate | Std. Error | z value | P value |
|---|---|---|---|---|
| Cyno / ZIKV - control == 0 | -1.32 | 0.80 | -1.643 | 0.10 |
| Squirrel / ZIKV - control == 0 | -0.45 | 0.31 | -1.430 | 0.15 |
| **Squirrel / DENV2 - control == 0** | -1.19 | 0.32 | -3.716 | **2.02e-04** |
| Cyno / control - DENV2 == 0 | 0.18 | 0.78 | 0.231 | 0.82 |
| Control / Cyno - Squirrel == 0 | 0.86 | 0.67 | 1.280 | 0.20 |
| **DENV2 / Cyno - Squirrel == 0** | 1.87 | 0.52 | 3.599 | **3.2e-04** |
| ZIKV / Cyno - Squirrel == 0 | -0.01 | 0.54 | -0.017 | 0.99 |

Table S.4: Comparison of mosquito engorgement on day 0, based on mosquito infection status and monkey species, for the selected model m1. For model description and selection, see Tables S.5, S.6.

| Model description | | Model name | AICc |
|---|---|---|---|
| Error distribution | Random effects | | |
| binomial | no | m0 | 315.3 |
| betabinomial | no | m1 | **284.4** |
| binomial | yes | m2 | 292.6 |
| betabinomial | yes | m3 | 286.9 |

Table S.5: Selection based on corrected Akaike Information Criterion (AICc) for models testing the impact of mosquito infection status on engorgement rates on day 0 of the experiments. Control and cofeeding mosquitoes were aggregated, per species. Random effects, when included, consisted in fitting an intercept per monkey.

| Likelihood ratio tests | | | | |
|---|---|---|---|---|
| | m0 | m1 | m2 | m3 |
| m0 | | 6.8 e-09 | 4.7 e-07 | |
| m1 | | | | 0.68 |
| m2 | | | | 3.8 e-03 |
| m3 | | | | |

Table S.6: Selection based on likelihood ratio tests between nested models. $P$ values are reported, and when they're $< 0.05$, the more complex model (ordered by increasing number) is selected. For model description see Table S.5.

### Feeding of uninfected mosquitoes on infected versus uninfected hosts: Experiment Days 1-21

#### Simple model

| Comparison | Estimate | Std. Error | z value | P value |
|---|---|---|---|---|
| **DENV2 / Cyno - Squirrel == 0** | 1.02 | 0.18 | 5.82 | **5.84e-09** |
| ZIKV / Cyno - Squirrel == 0 | 0.10 | 0.20 | 0.51 | 0.61 |
| **Control / Cyno - Squirrel == 0** | 0.74 | 0.26 | 2.86 | **0.0043** |
| Cyno / DENV2 - control == 0 | 0.18 | 0.22 | 0.82 | 0.41 |
| **Cyno / ZIKV - control == 0** | -0.81 | 0.24 | -3.36 | **0.00078** |
| Squirrel / DENV2 - control == 0 | -0.097 | 0.22 | -0.44 | 0.66 |
| Squirrel / ZIKV - control == 0 | -0.17 | 0.22 | -0.78 | 0.44 |

Table S.7: Comparison of mosquito engorgement on days post monkey infection, based on monkey infection status and monkey species, for the selected simple model m1. For model description and selection, see Tables S.8, S.9.

| Model description | | Model name | AICc |
|---|---|---|---|
| Error distribution | Random effects | | |
| binomial | no | m0 | 1685.7 |
| betabinomial | no | m1 | **1410.1** |
| binomial | yes | m2 | 1656.9 |
| betabinomial | yes | m3 | 1412.2 |

Table S.8: Selection based on corrected Akaike Information Criterion (AICc) for simple models testing the impact of monkey infection status on engorgement rates. When they were included, random effects consisted in fitting an intercept per monkey and per day.

| Likelihood ratio tests | | | | |
|---|---|---|---|---|
| | m0 | m1 | m2 | m3 |
| m0 | | < 2.2 e-16 | 2.7 e-08 | |
| m1 | | | | 1 |
| m2 | | | | < 2.2 e-16 |
| m3 | | | | |

Table S.9: Selection based on likelihood ratio tests between nested models. *P* values are reported, and when they're < 0.05, the more complex model (ordered by increasing number) is selected. For model description see Table S.8

.

#### Complete model

| | Estimate | Std. Error | z value | P value |
|---|---|---|---|---|
| Intercept* | -1.98 | 1.77 | -1.12 | 0.26 |
| viremia° | -0.045 | 0.057 | -0.79 | 0.43 |
| DENV2 - control | -0.045 | 0.19 | -0.24 | 0.81 |
| ZIKV - control | -0.28 | 0.18 | -1.56 | 0.12 |
| Male - Female | -0.071 | 0.18 | -0.39 | 0.70 |
| time of day† | -0.076 | 0.079 | -0.97 | 0.33 |
| Squirrel - Cyno | 1.45 | 1.91 | 0.76 | 0.45 |
| weight† | -0.39 | 1.84 | -0.21 | 0.83 |
| host body temperature† | 0.34 | 0.14 | 2.35 | 0.019 |
| weight * species | 0.22 | 1.79 | 0.13 | 0.90 |
| host body temperature * species | -0.36 | 0.21 | -1.67 | 0.095 |
| *Switching to Cyno as reference* | | | | |
| host body temperature† | -0.02141 | 0.16476 | -0.130 | 0.8966 |
| weight† | -0.16286 | 0.26219 | -0.621 | 0.5345 |

Table S.10: Results of the selected complete model m3. For model description and selection, see Tables S.11, S.12.
* : ref = DENV-2 infected male squirrel
° : effect of an increase of 1 $\log_{10}$ PFU/ml
† : effect of increase of 1 standard deviation (scaled variable)

| Model description | | Model name | AICc |
|---|---|---|---|
| Error distribution | Random effects | | |
| binomial | no | m0 | 1705.6 |
| betabinomial | no | m1 | 1424.4 |
| binomial | yes | m2 | 1608.4 |
| betabinomial | yes | m3 | **1420.3** |

Table S.11: Selection based on corrected Akaike Information Criterion (AICc) for complete models testing the impact of monkey infection status on engorgement rates. When they were included, random effects consisted in fitting an intercept per monkey and per day.

| Likelihood ratio tests | | | | |
|---|---|---|---|---|
| | m0 | m1 | m2 | m3 |
| m0 | | < 2.2 e-16 | < 2.2 e-16 | |
| m1 | | | | 0.015 |
| m2 | | | | < 2.2 e-16 |
| m3 | | | | |

Table S.12: Selection based on likelihood ratio tests between nested models. *P* values are reported, and when they're < 0.05, the more complex model (ordered by increasing number) is selected. For model description see Table S.11

.

#### S.2 Comparison with inter-individual variation in approach rate of free-living *Ae. albopictus*

Collection of approach data in Hernandez Acosta 2023 was done across three levels of urbanization, at the following collection windows: 6:00-7:00, 9:00-10:00, 12:00-13:00, 15:00-16:00, and 18:00-19:00 hours. Urbanization was measured by the Normalized Difference Built-up Index (NDBI) and classified using the Jenks natural breaks method into low (-1 to -0.467), medium (-0.468 to -0.098), and high (-0.099 to 1.000) NDBI value categories (Hendy et al. 2023).

From this larger dataset, we extracted the number of approaches per collector at times and at urbanization levels showing the highest number of female *Ae. albopictus* approaching, as well as the highest proportion of non-zero approach among attempts. These optimal conditions were observed for low and medium NDBI, and for collection windows 12:00-13:00 and 15:00-16:00.

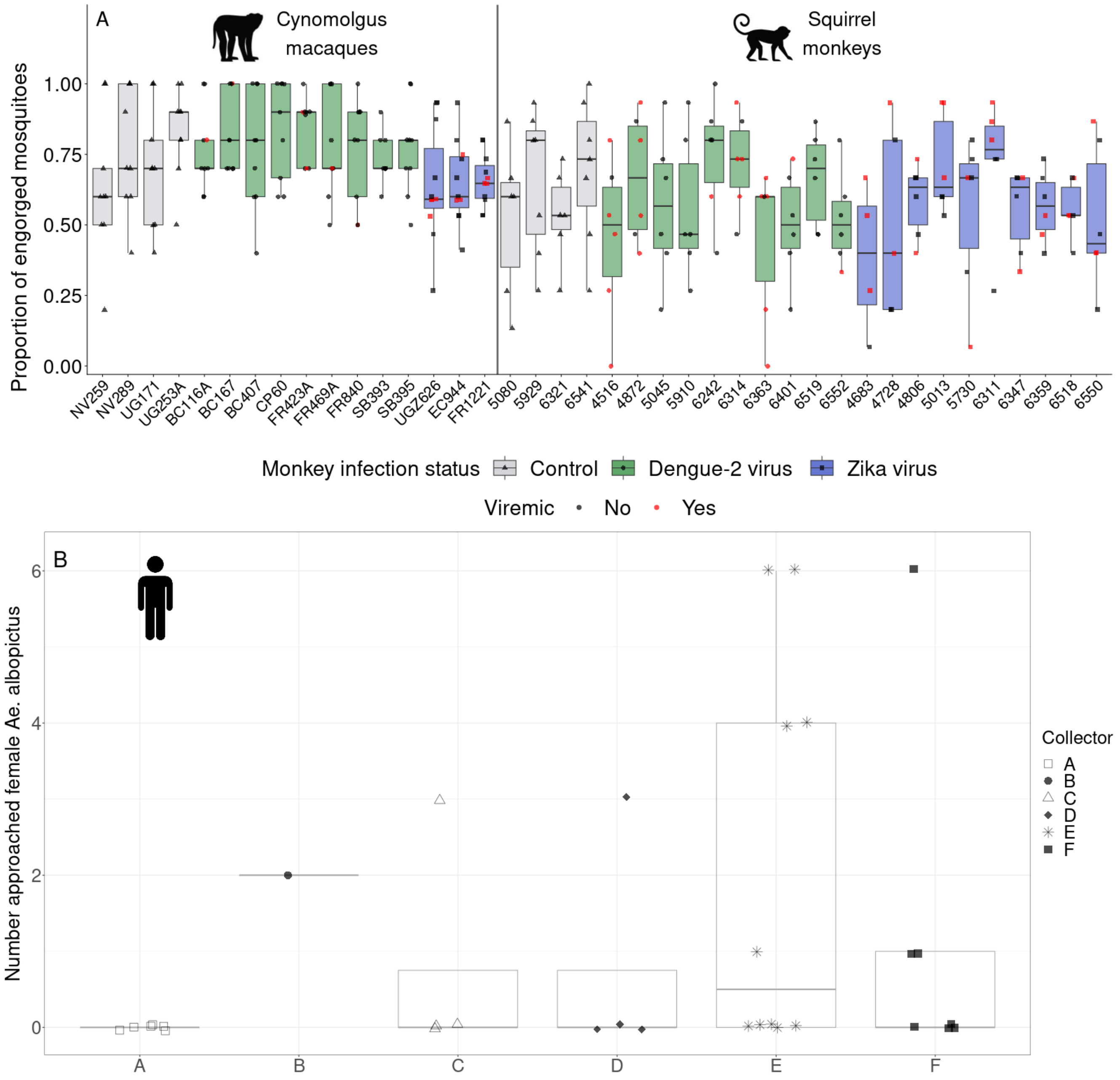


Figure S.2: Comparison of the variability of mosquito behavior between our experiment and field observations of free-living mosquitoes. **A** - Proportion of engorged *Ae. albopictus* on individual monkeys, depending on their infection status and species. One boxplot per monkey, colored by monkey infection status (grey : control, green : DENV-2 infected, blue : ZIKV-infected), with raw data (triangle : control, circle : DENV-2 infected, square : ZIKV-infected). Red points indicate engorgement on animals that were detectably viremic, or that resulted in transmission to mosquitoes. Data from day 28 excluded (see Methods). **B** - Approach data from Manaus, Brazil. Number of female *Ae. albopictus* which approached several human collectors, in optimum conditions defined as low and medium NDBI, and collection windows 12:00-13:00 and 15:00-16:00. For the coefficients of variation derived from this dataset, we excluded collector B, who only performed one attempt in these conditions, and we assigned a coefficient of variation of 0 to collector A who wasn't approached by any female *Ae. albopictus* in 6 attempts.

#### S.3 Cytokines

We selected between linear and generalized additive models using AICc. Competing models were fitted using maximum likelihood, for each monkey species and each cytokine separately. Monkey ID was used as a random effect (intercept). Day was not used as it did not improve AICc. In the additive model, the number of knots was constrained to 4 for fixed effects and to 2 for random intercepts, to avoid over-fitting. All concentrations below the limit of detection (LOD) were excluded from analysis initially, for all models. This substantially improved the residuals of the models, which were checked using the R package DHARMa. For each model type, we checked that significant results remained as such after using the Benjamini-Hochberg correction for $P$ values, with a false discovery rate (false positives / (false positives + true positives)) of 5% (Benjamini and Hochberg 1995), but we report the initial $P$ values in the results section and in the present document. For the selected model (linear or additive), if a result stood out after $P$ value correction, the same model including measures below LOD (in that case, fixed at LOD) was run and compared.

All results are presented in Table S.9 (separate file). Below are some details for the cytokines showing significant associations with bite exposure.
For **TGF-β**, a generalized additive mixed effect model was selected, estimating linear effects of both short-term and long-term bite exposure, and splines to fit individual intercepts. The concentration of TGF-β was negatively associated with the cumulative number of bites received in the last 7 days (-0.0036 [-0.0060 ; -0.0013] $\log_{10}$ pg/µl per additional bite, p = 0.0054).
For **MCP-1**, a generalized additive mixed effect model was selected, estimating linear effects of both short-term and long-term bite exposure, and splines to fit individual intercepts. The concentration of MCP-1 was negatively associated with the cumulative number of bites received in the last 7 days (-0.0039 [-0.0064 ; -0.0013] $\log_{10}$ pg/µl per additional bite, p = 0.0058).

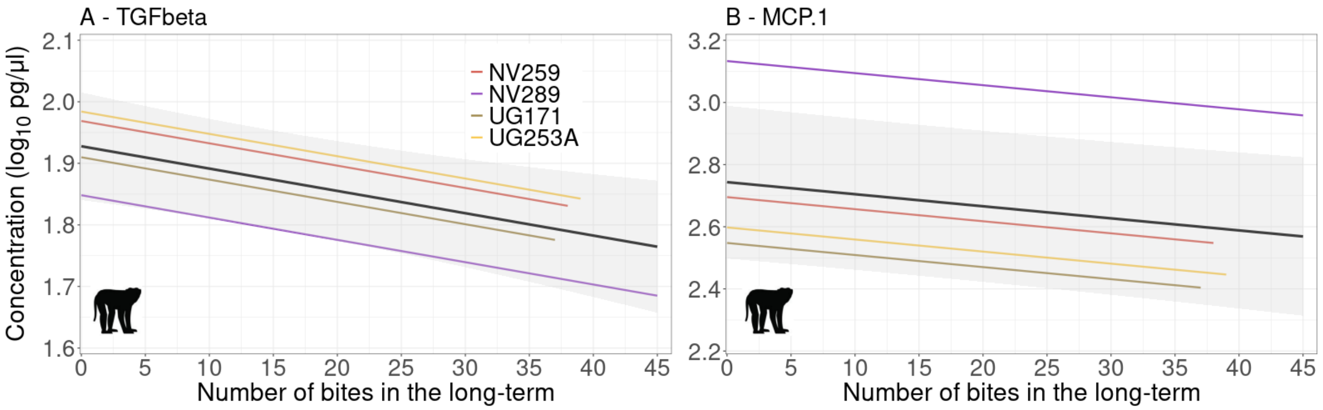


Figure S.3: In control cynomolgus macaques, concentrations of cytokines TGF-β (A) and MCP-1 (B) are significantly impacted by the number of uninfected mosquitoes which engorged in the long-term (last 7 days). Results from generalized additive mixed effect models, estimating an intercept per monkey. Thick black lines and shading show the population trends with uncertainty. Colored lines are individual fits. Fits are computed with the short-term exposure variable (previous day) fixed at 0, which is why datapoints are not plotted (short-term and long-term variables vary concomitantly). Note that y-axis scales differ between cytokines.

For **EGF**, a generalized additive mixed effect model was selected, estimating linear effects of both short-term and long-term bite exposure, and splines to fit individual intercepts. EGF concentrations increased by 0.02 [0.01 ; 0.03] log10 pg/µl per additional bite the day prior (p = 2.15e-4, see Figure 6A of main text), but decreased by 0.006 [-0.008 ; -0.004] log10 pg/µl per additional bite in the last seven days (p = 8e-6, see Figure 6C in main text).
For **MIF** a generalized additive mixed effect model was selected, estimating a linear effect of long-term bite exposure, and splines for short-term bite exposure and individual intercepts. The relationship with short-term exposure was non-linearly positive, with a saturation after about 6 bites the day prior (p = 0.005, see Figure 6B in main text). MIf concentrations decreased by 0.010 [-0.014 ; -0.005] log10 pg/µl per additional bite in the last seven days (p = 2.3e-4, see Figure 6D in main text).
